# How human-derived brain organoids are built differently from brain organoids derived of genetically-close relatives: A multi-scale hypothesis

**DOI:** 10.1101/2023.05.25.542171

**Authors:** Tao Zhang, Sarthak Gupta, Madeline A. Lancaster, J. M. Schwarz

## Abstract

How genes affect tissue scale organization remains a longstanding biological puzzle. As experimental efforts are underway to solve this puzzle via quantification of gene expression, chromatin organization, cellular structure, and tissue structure, computational modeling efforts remain far behind. To help accelerate the computational modeling efforts, we review two recent publications, the first on a cellular-based model for tissues and the second on a model of a cell nucleus that includes a lamina shell and chromatin. We then address how the two models can be combined to ultimately test multiscale hypotheses linking the chromatin scale and the tissue scale. To be concrete, we turn to an *in vitro* system for the brain known as a brain organoid. We provide a multiscale hypothesis to distinguish structural differences between brain organoids built from induced-pluripotent human stem cells and from induced-pluripotent gorilla and chimpanzee stem cells. Recent experiments discover that a cell fate transition from neuroepithelial cells to radial glial cells includes a new intermediate state that is delayed in human-derived brain organoids as compared to their genetically-close relatives, which significantly narrows and lengthens the cells on the apical side [1]. Additional experiments revealed that the protein ZEB2 plays a major role in the emergence of this new intermediate state with ZEB2 mRNA levels peaking at the onset of the emergence [1]. We postulate that the enhancement of ZEB2 expression driving this intermediate state is potentially due to chromatin reorganization. More precisely, there exists critical strain triggering the reorganization that is higher for human-derived stem cells, thereby resulting in a delay. Such a hypothesis can readily be tested experimentally within individual cells and within brain organoids as well as computationally to help work towards solving the gene-to-tissue organization puzzle.

## I. INTRODUCTION

Genetic mutations can indeed impact tissue scale organization. For instance, there is plentiful experimental evidence that changes in gene expression can affect the foliated structure of a developing brain [2–5]. To be even more specific, mutations of the LIS1 gene result in lissencephaly, or a smooth brain [6, 7]. And while the connection between genes and tissue scale organization is highly complex, of which many puzzle pieces remain unknown, experimental efforts are underway to find the missing puzzle pieces. Recent experimental progress on linking the chromatin scale with the tissue scale is now emerging with, for example, the finding that mechanical straining a tissue leads to the loss of heterochromatin to give rise to cell nuclear softening [8]. Structural measurements at the cell and tissue scale have long been standard in biology. Measurements of the spatial organization of chromatin in cells using chromatin conformation capture techniques are now also well-established [9–16]. Other methodologies acquiring additional information about chromatin architecture include immunoGAM [17] and spatially resolving chromatin modifications [18] will also help put together this puzzle. Given these experimental developments, one wonders how the current state of computational modeling can also help solve this puzzle. This manuscript gives a roadmap on how to begin to build minimal, multiscale computational models to help solve this puzzle.

Given some recent, intriguing experimental results on brain organoids [1], we will use these results as a guide. So we now zero in on the brain. How does the brain work? One could argue that to understand how something works, one must be able to build it. As a brain is being built, it is composed of living, multiscale matter capable of emergent forms of mechanical, chemical, and electrical functionality at the genome scale, the cell nucleus scale, the cellular scale, and/or the tissue scale in a nested structure with interplay between the different scales. While such a multiscale materials neuroscience viewpoint may seem obvious-but-unwieldy to many, given the theoretical and experimental techniques that have evolved over the past few decades, we are now on the cusp of being able to develop quantitative predictions based on this viewpoint for the brain structure-function relationship, and, more importantly, to test them using brain organoids [19–21]—an *in vitro* realization of a developing brain. See Figure 1.

**FIG. 1.**
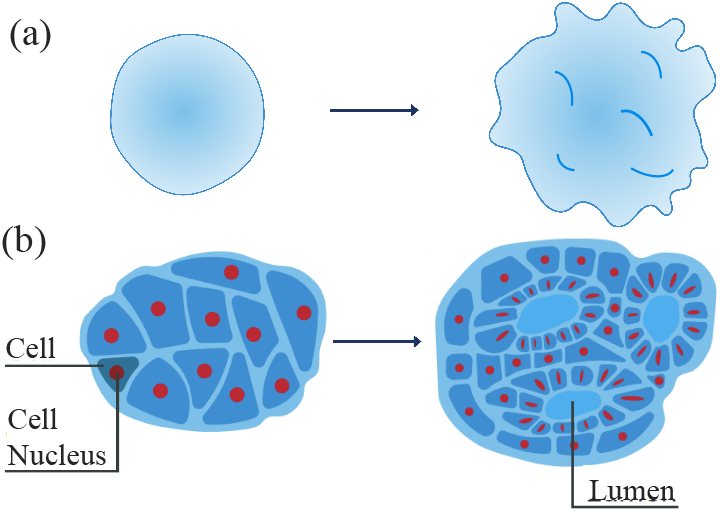
Brain organoid schematics. (a) Schematic of a brain organoid at an earlier time and at a later time. (b) Schematic of a cross-section of (a) with more cellular detail.

To initiate this multiscale materials neuroscience view-point in a computational setting, here, we review two recent computational models, one at the tissue scale developed to quantify the structure and mechanics of an confluent cellular collective, including brain organoids early in development, and a second at the cell nucleus scale developed to quantify correlated chromatin motion and cell nuclear shape fluctuations [22, 23]. After reviewing these two models, we will then demonstrate how they can be coupled to continue to probe a key brain organoid structure question that has been recently asked and begun to be answered experimentally: How does the development of the structure of human-derived brain organoids differ from their closest genetic relatives, namely chimpanzee-derived and gorilla-derived brain organoids? More specifically, we will briefly review the experiments, formulate a testable, multiscale hypothesis and then demonstrate, in principle, how it can be first tested computationally to determine its feasibility. In other words, we provide a roadmap for a direction of next-generation multi-scale, cellular-based computational models that are minimal— minimal in the sense that complexity emerges from simplicity as opposed to complexity emerging from complexity. They can also be falsified by experiments as they yield predictions about organoid shape and rheology, cell shape and rheology and cell nucleus shape and rheology and chromatin structure, ultimately. With falsification, comes progress. Without falsification, one can travel down a wrong road for a lengthy amount of time. See Figure 2 for an overview of the computational frame-work that we will elucidate as the manuscript unfolds.

**FIG. 2.**
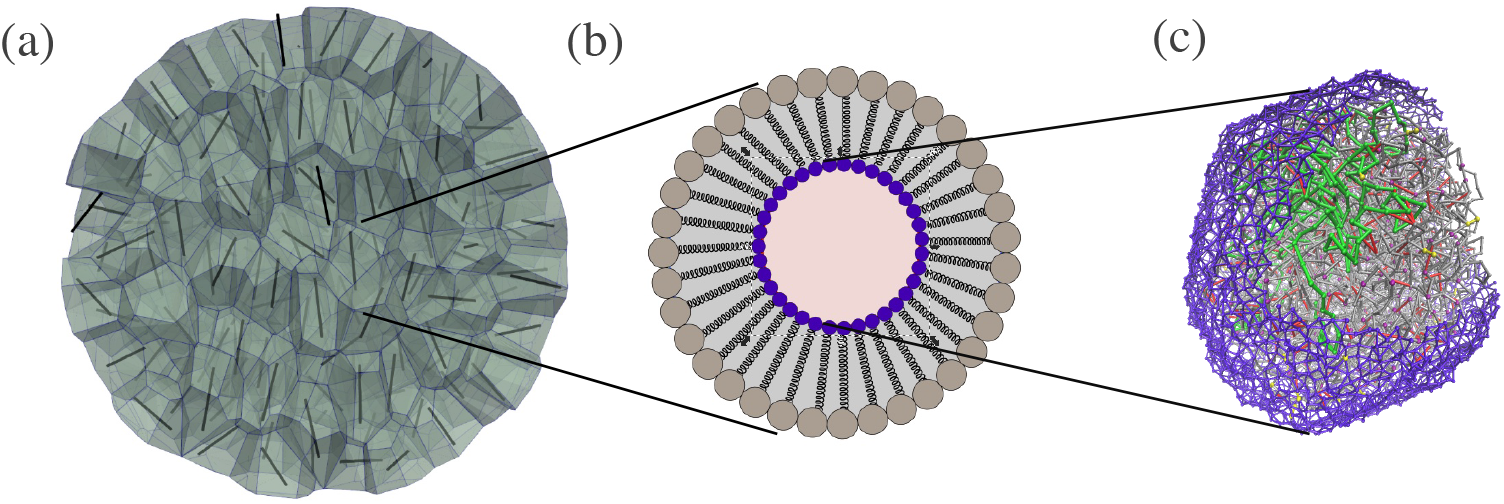
A computational approach to multiscale modeling of tissue structure: (a) A representative organoid based on a three-dimensional vertex model in which cells are represented as deformable polyhedrons and there are no gaps between the cells. The black rods denote the long axis of the polyhedron as determined by a fit to a minimal volume ellipsoid. (b) A schematic of a two-dimensional cross-section of a cell that includes the acto-myosin cortex (outer ring of springs), the lamina shell (inner ring of springs), and the bulk cytoskeleton, including vimentin (the springs connecting the inner and outer ring of springs). (c) A deformable lamina shell cell nuclei (purple) containing chromatin (grey). A portion of the chromatin is colored in green to highlight its configuration.

## II. A CELLULAR-BASED COMPUTATIONAL MODEL FOR ORGANOIDS

Let us begin with a cellular-based, computational model for an organoid that is rooted in earlier work [24–31]. With such a model, we can track changes in cell shape, which can potentially give rise to changes in nuclear shape such that changes in nuclear shape can potentially lead to changes in chromatin organization. To this end, let us review recent construction of a what is called a three-dimensional vertex model with boundaries [22].

Cells are biomechanical and biochemical learning machines that are not in equilibrium, i.e., they are driven by internal, or active forces. The biomechanics of the organoid, which is a collection of cells, is given by the energy functional:

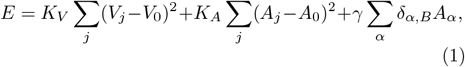

where *A*_*j*_ denotes the *j*th cell total area, the *j*th cell volume is denoted by *V*_*j*_, and *α* labels the faces of the cells, with *δ*_*α,B*_ = 0 if a cellular face is not at the boundary B of the collective and 1 otherwise. Given the quadratic penalty from deviating from a cell’s preferred volume and area, *K*_*V*_ and *K*_*A*_ are volume and area stiffnesses, respectively. Physically, the volume term represents the bulk elasticity of the cell with *V*_0_ denoting a target volume. If the area term is expanded to contain a quadratic contribution, a linear contribution, and a constant contribution, it has typically been argued that the quadratic term represents the contractility of the acto-myosin cortex and the linear term, whose coefficient is −2*K*_*A*_*A*_0_, is determined by a competition between cell-cell adhesion and cortical contractility [26, 30]. The latter dominates at negative values, corresponding to smaller values of *A*_0_, and the former dominates at positive values, corresponding to larger values of *A*_0_.

Indeed, cell-cell adhesion and contractility are coupled [32]. For instance, knocking out E-cadherin in keratinocytes, effectively changes the contractility [31]. Given this intricate coupling, it may be difficult to tease out the competition. Moreover, the finding in two-dimensional vertex models of a rigidity transition as the target perimeter is increased then leads to the interpretation that unjamming, or fluidity, is given by an increase in cell-cell adhesion, which appears to be counterintuitive [30]. As for an alternative interpretation, by adding a constant to the energy, which does not influence the forces, again, the energy can be written in the above quadratic form. Since cell we cannot tune cell-cell adhesion independently of cortical contractility, we posit that the target area is simply a measure of the isotropy of cortical contractility, assuming that curvature changes in the cells remain at scales much smaller than the inverse of a typical edge length. It is the cell-cell adhesion that is bootstrapped to the cortical contractility as cell faces are always shared. To be specific, the larger the target area, the less isotropically contractile the cell is, and vice versa. The less isotropically contractile a cell is, the more it can explore other shapes to be able to move past each other in an energy barrier-free manner resulting in fluidity. Additional terms linear in the area for specific cell faces complexify the notion of isotropic contractility.

As for the linear area term in Eq. (1), the cells at the boundary of the organoid, there is an additional surface tension term for faces interacting with the “vacuum” consisting of empty cells. These empty cell do not exert forces on the cells but allow the cells on the surface of the organoid to relax. We also restrict cells from separating from the organoid, at least at this time. One can nondimensionalize any length *l* in the simulation with 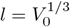. As one does so, an important parameter in these models is the dimensionless shape index 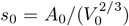, which is related to the target area. A regular tetrahedron has a dimensionless shape index of *s*_0_ ≈ 7.2, for example.

We have addressed the biomechanical aspect of cells with biochemical aspects indirectly encoded into the model parameters. We must also account for their dynamics. Cells can move past each other even when there are no gaps between them. In two dimensions, such movements are known as T1 events. Understanding these events are key to understanding the rigidity transition in two dimensions [30]. In three dimensions, such movements are known as reconnection events. Prior work has developed an algorithm for such reconnection events focusing on edges becoming triangles and vice versa that may occur for edges below a threshold length *l*_*th*_ for a fixed topology [28]. Specifically, each vertex has four neighboring vertices and shares four neighbor cells. Each edge shares three cells and each face/polygon shares two cells. Our modeling builds on that key work [28].

In addition to reconnection events, there is an underlying Brownian dynamics for each vertex, hence, the term “vertex model”. In other words, the equation of motion for the position **r**_*I*_ of a single vertex *I* is

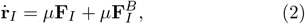

with **F**_*I*_ and 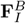 denoting the conservative force and the random force due to active fluctuations on the *Ith* vertex respectively. The force **F**_*I*_ is determined from both the area and volume energetic constraints and, hence, includes cell-cell interactions. Moreover, each vertex performs a random walk with an effective diffusion coefficient of *μk*_*B*_*T*, where *T* is an effective temperature. Unless otherwise specified, the mobility *μ* = 1. Finally, the Euler-Maruyama integration method is used to update the position of each vertex.

As we have constructed a three-dimensional vertex model with a quadratic energy functional in terms of a target surface area and a target volume, we then look a rigidity transition in the bulk system with periodic boundary conditions from a fluid-like state and a solid-like state. Such a rigidity transition occurs in two dimensions as the target shape index decreases [30]. By measuring a neighbor overlap function that keeps track of whether or not a cell loses 2 or more of its neighbors over time, as would occur if cells are readily moving throughout the system, we determined the transition location to occur at a target shape index of 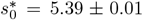, for the system sizes and effective temperatures studied. In Figure 3(a) and (b) we show trajectories of cells to give an indication of the rigidity transition. This transition location is slightly lower than the location of the rigidity transition observed in a three-dimensional Voronoi version, where the degrees of freedom are assigned to the cell centers, using energy minimization [33]. The fluidization is driven by a decrease in isotropic contractility and allows the emergence of new cellular faces in three dimensions as the polyhedrons are able to explore more configurations. Moreover, the decrease in isotropic contractility may lead to an increases in anisotropic contractility via stress fibers may ultimately drive cell motion. Such an effect in single cells has been recently emphasized [34] and is likely to occur at the multi-cellular level [35].

**FIG. 3.**
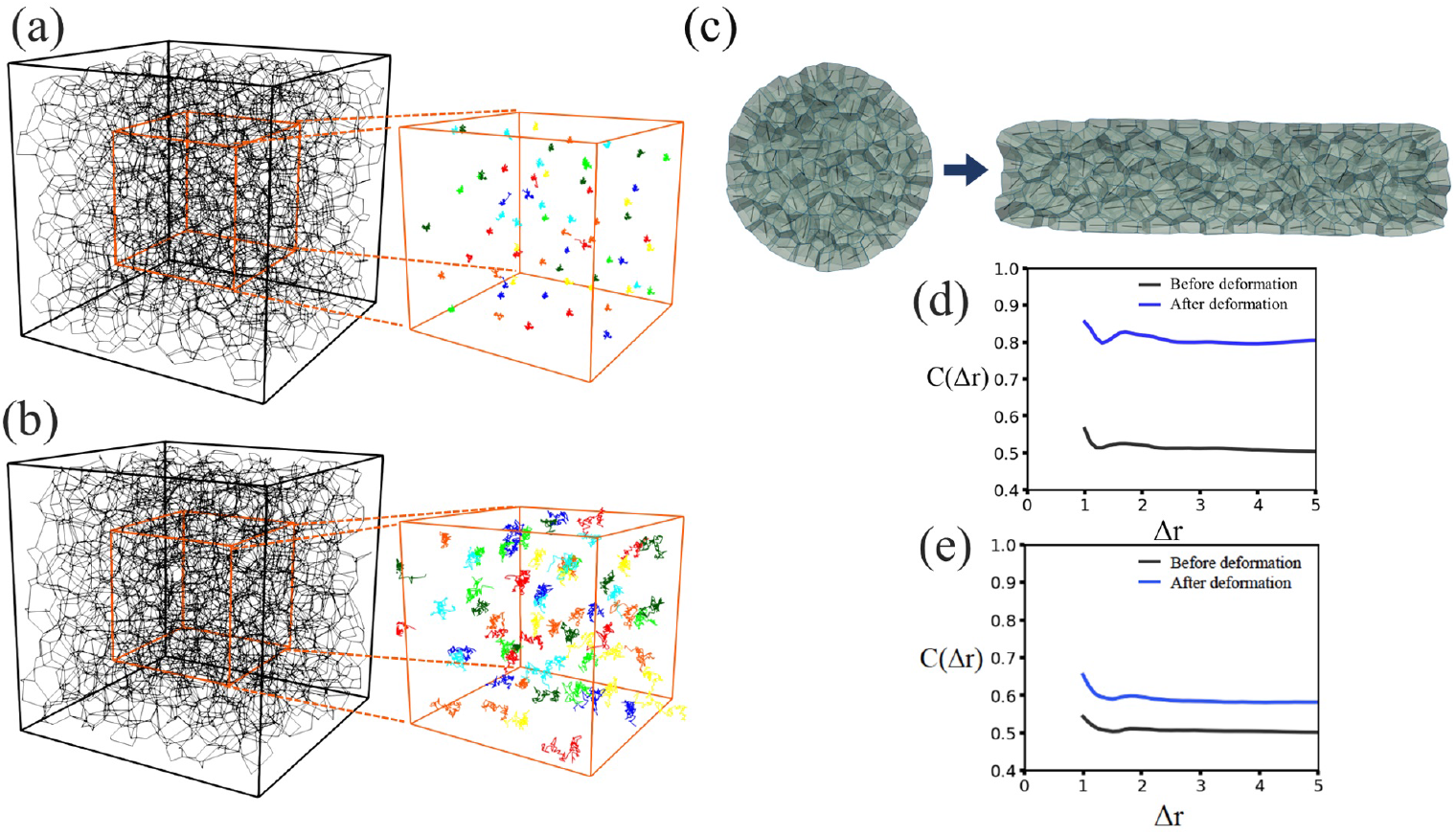
A three-dimensional vertex model: (a) The edges of the cells are depicted in black, with an edge loop representing a cell face. The target shape index, or *s*_0_ = *A*_0_*/*(*V*_0_)^2*/*3^ = 5.3, with *A* denoting surface area of a cell and *V* denoting its volume. Periodic boundary conditions are implemented. For a subset of the system, the trajectories of the center-of-mass of the cells is plotted over the time. (b) Same as (a) except with *s*_0_ = 5.6. (c) Cross-sectional snapshots of the organoid before and after the lateral extensile deformation, where the black rods indicate the long-axis of the cell, or cell orientation, as determined by a fit to a minimal volume ellipsoid. (d) The cell-cell orientation correlation function *C*(Δ*r*) as a function of the distance Δ*r* between the centers of two cells before and after the lateral extensile deformation for cells in the bulk with *s*_0_ = 5.6; (e) Same as (d) though for cells on the boundary, again, with *s*_0_ = 5.6. The images are reprinted from Tao Zhang and J. M. Schwarz, *Phys. Rev. Res*. **4**, 043148 (2022).

Given the existence of a rigidity transition, it is natural to ask how does the transition affect the system’s response to deformations, such as lateral and in-plane radial extension. We indeed observe vestiges of the rigidity transition in the organoid case. More specifically, for both lateral and in-plane radial extension, we observe larger changes in the structure of the cells before and after the deformation for the fluid-like organoid as compared to the solid-like organoid. What do we mean by structure? For lateral extension, for instance, the boundary cells align along the direction of the deformation, whereas the cells in the bulk do not. In other words, the bulk cells resemble the cells in the bulk system with periodic boundary conditions. We quantify this alignment by measuring the cell-cell orientation correlation function for the long axis of each cell as a function of distance between the centers of cells, or *C*(Δ*r*). See Figures 3(c)-(e). For cells not aligned along their long axis, 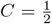, while for cells aligned along their long axis *C* = 1.

This stark difference between the bulk and boundary cells is a phenomenon that cannot be readily captured in a continuum model. And while there are other ways to depict three-dimensional cellular collectives, such as cellular Potts model [36] or a three-dimensional Voronoi model [33, 37, 38], the three-dimensional vertex model is our model of choice as it contains detailed information about cell shape with the degrees of freedom associated with an acto-myosin cortex and shared cell faces. The topologically-protected interior is a consequence of the absence of any interfacial surface tension at the inner faces of the boundary cells. While the boundary cells appear columnar, the lack of interfacial surface tension on the inner faces does not imply right prisms in which the boundary face and the opposite, inner face are parallel. As long as these inner faces are not constrained to be parallel to the boundary faces, the cells do not retain memory of the smooth boundary. In other words, with this featur,e the cells can take on a more varied zoology of shapes beyond right prisms observed a two-layered, two cell-type Voronoi model with interfacial surface tension between two cell types [37].

The single-cell-layer thick boundary effects provide a mechanism for patterning with a confluent cellular collective in which its interior remains topologically-protected from deformations at the boundary. For instance, quasi-two-dimensional brain organoids have a cortex that is approximately one-cell layer thick [39]. In terms of timescales, at least over time scales larger than the time scale for cellular rearrangements, the boundary cells are indeed insulating the bulk cells from the boundary deformation as the confluent cellular collective deforms. Over shorter times, the shape of the cells mimic the boundary deformation, just as an elastic solid. One can therefore observe in a developmental system, for instance, different regimes of deformations at the cell scale in Drosophila epithelial morphogenesis [40].

While the vertex model is a cellular-based model, there is no explicit cellular contents except for effectively an acto-myosin cortex surrounding an effectively incompressible fluid with adhesion molecules that maintain the perfectly zipped-up cell surfaces. Such a model is sufficient to probe many questions, however, there exist many sets of questions where more explicit cellular content is required, such as an explicit cell nucleus. With the cell nucleus being the most rigid structure in the cell [41], presumably, for more extreme mechanical perturbations of tissues, cell nuclei set limits on the deformability of the tissue, just as it has been shown that cell nuclei set limits on individual cell motility [42–44]. Let us, therefore, discuss cell nuclei in more detail.

## III. A COMPUTATIONAL MODEL OF DEFORMABLE CELL NUCLEI

The cell nucleus houses the genome, or the material containing instructions for building the proteins that a cell needs to function. For humans and other genetically-close relatives, this material is ∼ 1 meter of DNA. Using proteins to form chromatin, the DNA is packaged across multiple spatial scales to fit inside an ∼ 10 *μ*m nucleus [45]. In addition, chromatin is highly dynamic; for instance, correlated motion of micron-scale genomic regions over timescales of tens of seconds has been observed in mammalian cell nuclei [46–50]. This correlated motion diminishes both in the absence of ATP, the fuel for many molecular motors, and by inhibition of the transcription motor RNA polymerase II, suggesting that motor activity plays a key role [46, 47]. These dynamics occur within the confinement of the cell nucleus, which is enclosed by a double membrane and 10-30-nm thick filamentous layer of lamin intermediate filaments to form a lamina shell [51–53]. The lamina shell is deformable and, as such, one can quantify its shape fluctuations. Specifically, depletion of ATP diminishes the magnitude of the shape fluctuations, as does the inhibition of RNA polymerase II transcription activity [54].

Chromatin and the lamina shell interact directly via lamina-associated domains (LADs) [55, 56] and indirectly through various proteins [57–59]. Therefore, the spatiotemporal properties of chromatin can potentially influence shape of the lamina shell and vice versa as the two components are coupled. Indeed, studies have found that depleting linkages between chromatin and the nuclear lamina, or membrane, results in more deformable nuclei [60, 61], enhanced curvature fluctuations [62], and/or abnormal nuclear shapes [63]. Another recent study suggests that inhibiting motor activity diminishes nuclear bleb formation [64]. Moreover, depletion of lamin A in several human cell lines leads to increased diffusion of chromatin, suggesting that chromatin dynamics is also affected by linkages to the lamina [65]. Together, these experiments demonstrate the critical role of chromatin and its interplay with the lamina shell in determining nuclear shape.

To quantify chromatin dynamics *and* nuclear shape, we constructed a chromatin-lamina system with the chromatin modeled as an *active* Rouse chain and the lamina as an elastic, polymeric shell with linkages between the chain and the shell. We also included chromatin crosslinks, which may be a consequence of motors forming droplets [66] and/or complexes [67], as well as chromatin binding by proteins, such as heterochromatin protein I (HP1) [68]. Recent rheological measurements of the nucleus support the notion of chromatin crosslinks [69, 70], as does indirect evidence from chromosome conformation capture (Hi-C) [71]. Unlike previous chain-enclosed-by-a-deformable-shell models [62, 69, 70], our model has motor activity. We implemented the simplest type of motor, namely extensile and contractile monopoles that act non-reciprocally on the chromatin.

To be even more specific, interphase chromatin is modeled as a Rouse chain consisting of *N* monomers with radius *r*_*c*_ connected by Hookean springs with spring constant *K*. We include excluded volume interactions with a repulsive, soft-core potential between any two monomers, *ij*, and a distance between their centers denoted as 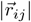, as given by

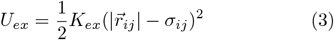

for 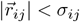, where 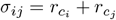, and zero otherwise. We include *N*_*C*_ crosslinks between chromatin monomers by introducing a spring between different parts of the chain with the same spring constant as along the chain. In addition to Gaussian fluctuations, we also allow for explicit motor activity along the chain. To do so, we assign some number, *N*_*m*_, of chain monomers to be active. An active monomer has motor strength *M* and exerts force **F**_*a*_ on monomers within a fixed range. Such a force may be attractive or “contractile,” drawing in chain monomers, or alternatively, repulsive or “extensile,” pushing them away. Since motors *in vivo* are dynamic, turning off after some characteristic time, we include a turnover timescale for the motor monomers *τ*_*m*_, after which a motor moves to another position on the chromatin. See Figure 2(c).

The lamina is modeled as a layer of *M* monomers connected by springs with the same radii and spring constants as the chain monomers and an average coordination number *z* ≈ 4.5, as supported by previous modeling [62, 69, 70] and imaging experiments [51–53]. We modeled the chromatin-lamina linkages as *N*_*L*_ permanent springs with stiffness *K* between shell monomers and chain monomers (Fig. 2(c)). There is an additional softcore repulsion between monomers making up the lamina shell to include excluded volume. See, again, Figure 2(c).

The system, as is the case for the three-dimensional vertex model, evolves via Brownian dynamics, obeying the overdamped equation of motion:

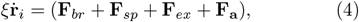

where **F**_*br*_ denotes the Brownian/Gaussian force, **F**_*sp*_ denotes the harmonic forces due to chain springs, chromatin crosslink springs, and chromatin-lamina linkage springs, and **F**_*ex*_ denotes the force due to excluded volume. In addition to the deformable shell, we also simulate a hard shell by freezing out the motion of the shell monomers.

We then studied the steady-state properties of this composite chromatin-lamina system in the presence of activity, crosslinking, rigidity of the lamina shell, and number of linkages between chromatin and the lamina. As for the range of number of crosslinks studied, we restricted the range to be such that chromatin exhibited anomalous diffusion, as observed experimentally [65], with a crossover to a smaller anomalous exponent driven by the crosslinking [72], since as the number of crosslinks increases, the chromatin eventually gels. For a deformable, lamina shell and in the presence of linkages, crosslinks, and motors, our model captures the correlated chromatin motion on the scale of the nucleus over various time windows. See Figures 4(a)-(g). The correlated motion occurs for both types of motors—-contractile or extensile—-and the deformability of the shell also plays a role. More precisely, for rigid lamina shells, the chromatin correlations are significantly diminished.

**FIG. 4.**
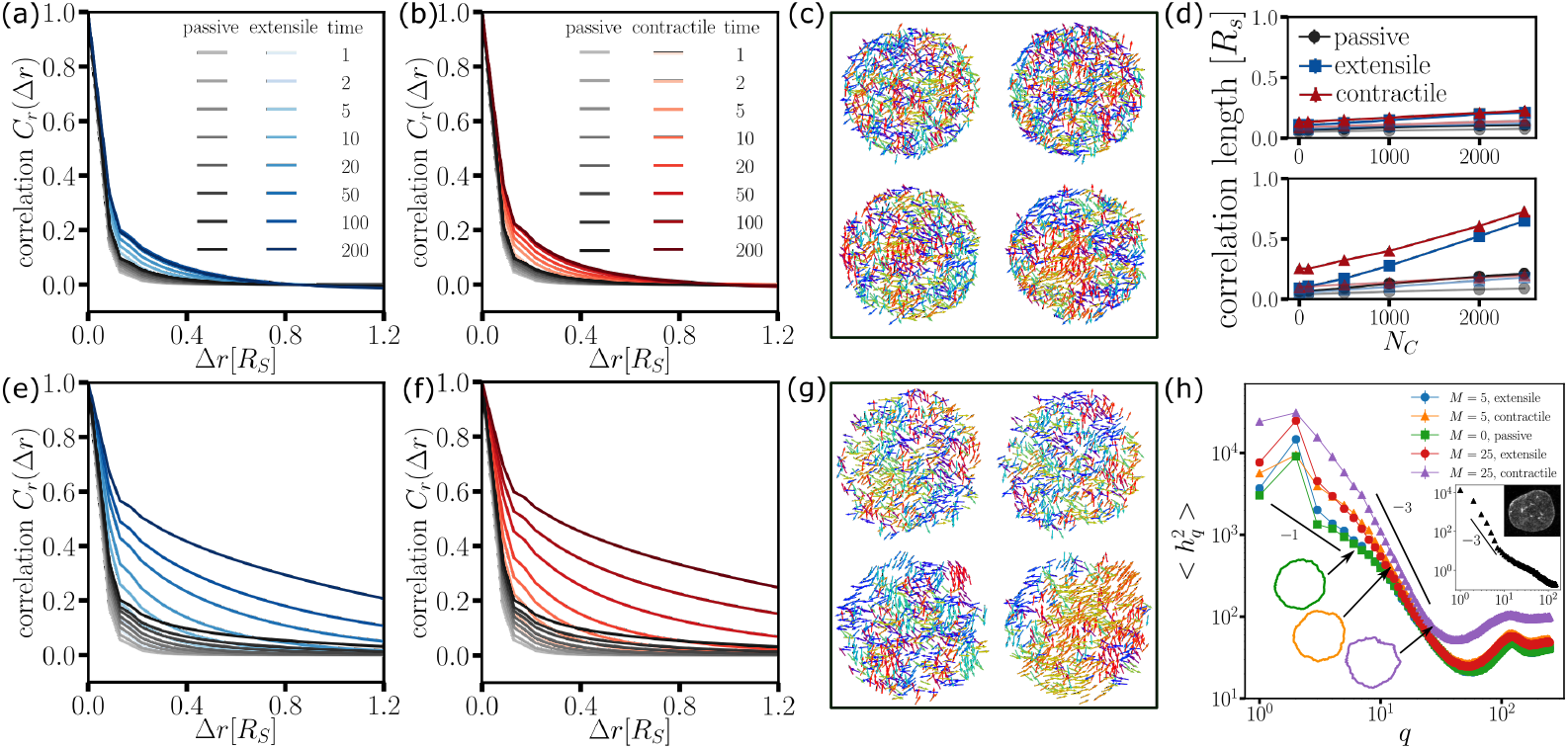
A cell nucleus model: (a) To look for correlated chromatin motion over some time scale, the spatial autocorrelation function of chromatin monomers *C*_*r*_ (Δ*r*, Δ*τ*) is plotted as a function of Δ*r* for different time windows and for passive (grey) and extensile motors (blue) at different time lags, Δ*τ*, for a hard lamina shell, while (b) s
hows the passive and contractile motors (red) case. (c) Two-dimensional vector fields for Δ*τ* = 5 (left), 50 (right) for the passive case (top) and the contractile case (bottom) from (b). (d) The correlation length, extracted from a phenomenological fit, as a function of the number of chromatin crosslinks *N*_*C*_ for the two time lags in (c) for the hard lamina shell (top) and soft lamina shell (bottom). (e∼g): The bottom row shows the same as the top row, but with a soft lamina shell. (h) Power spectrum of the shape fluctuations with *N*_*L*_ = 50 and *N*_*C*_ = 2500 for the passive and both active cases. Different motor strengths *M* are shown. The insets shows experimental data from mouse embryonic fibroblasts with an image of a nucleus with lamin A/C stained. The images are preprinted from Kuang Liu, Alison E. Patteson, Edward J. Banigan, and J. M. Schwarz, *Phys. Rev. Lett*. **126**, 158101 (2021).

**FIG. 5.**
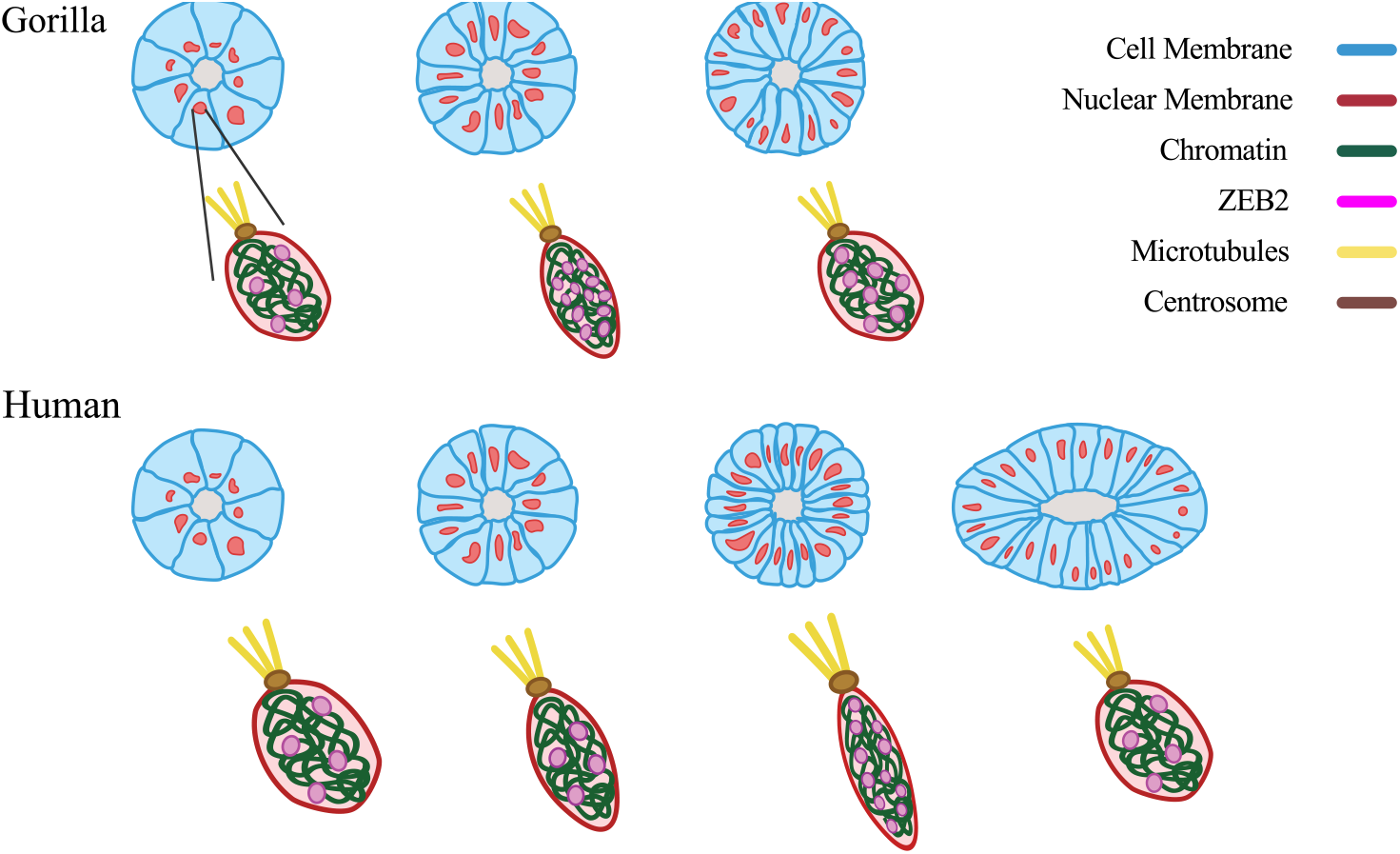
A multiscale hypothesis: The top row is a sketch of the time evolution of the cross-section of one cortex-lumen structure in a gorilla-derived brain organoid. The bottom row depicts the cross-section of the same structure, but in a human-derived brain organoid. According to our multiscale hypothesis, there is a critical strain on a cell nucleus to help initiate upregulation of ZEB2. The critical strain is larger for human-derived pluripotent stem cells as compared to gorilla-, or chimpanzee-derived pluripotent stem cells, resulting in a delay in the human-derived brain organoids. Note that ZEB2 can take on multiple roles, including inhibiting BMP-SMAD signaling by inhibiting BMP4 transcription to disrupt cell-cell junction formation and regulating the production of the microtubule-centrosome binding protein ninein. Graphics credit: Savana Swoger.

What is the role of deformability of the lamina shell? As motors amplify the motion of the chromatin, there are more distortions of the lamina shell. Since the lamina shell is elastic, these distortions are long-ranged, as evidenced by the broad power spectral density fluctuations. The long-ranged distortions in the lamina shell then in interacts back on the chromatin to result in an effectively long-range chromatin interaction, as mediated by the lamina shell. In other words, even though the chromatin is liquid-like, interaction with the lamina shell generates system-sized correlations along the chromatin chain. Moreover, as expected, increasing the number of crosslinks increases the chromatin correlation length with the absence of crosslinks leading to significantly lower correlations. Therefore, the chromatin crosslinks are required to generate a small coherent “patch” of chromatin in order to deform the shell sufficiently more than the fluctuations, as the motors act only on the chromatin. Also, as the number of motors decreases, the correlation length decreases as well, as expected.

As for understanding cell nuclear shape fluctuations, the power spectral density (PSD), or the magnitude of Fourier Transform of the height difference from a simple shape as a function of wavenumber *q*, of a two-dimensional cross-section exhibits power-law scaling. See Figure 4(g). For smaller wavenumbers, the power-law scaling is approximately *q*^*−*1^. For larger wavenumbers, the power-law scaling is *q*^*−*3^. Such scaling is expected for a fluctuating elastic shell with stretching and bending energetic costs with the latter generated by the topological constraint of the spring network forming a spherical shell [73, 74]. In addition, the amplitude of the PSD increases with increasing motor strength, for example. Notably, the contractile motor case exhibits more dramatic changes in the shape fluctuations as a function of wavenumber as compared to the extensile case. This finding is driven more anomalous density fluctuations for the contractile motor case in comparison to the extensile motor case. In the latter case, the extensile motor forces do not compete with the soft-core repulsion forces, while in the former case, the two forces do compete leading to more complex, local flows and, hence, density fluctuations that are more broad, or anomalous, in the contractile case.

Our short-ranged, active chromatin tethered to a deformable lamina shell model should be contrasted with an earlier confined, active Rouse chain interacting with a solvent via long-range hydrodynamics confined to a rigid shell with no-slip boundary conditions [48]. While both models generate correlated chromatin dynamics, with the long-range hydrodynamics model, such correlations are generated only with extensile motors that drive local nematic ordering of the chromatin chain [48]. Given this strong reliance of the long-range hydrodynamics model on extensile motors, it is important to experimentally test this prediction as well as look for local nematic ordering. Moreover, our correlation lengths are significantly larger than those obtained in a confined active, heteropolymer confined to a rigid shell [75]. Activity in the heteropolymer model is incorporated as extra-strong thermal noise such that the correlation length decreases at longer time windows as compared to the passive case. This decrease contrasts with our results and with experiments [47]. Future experiments are most certainly required to potentially distinguish the various proposed mechanisms.

In sum, the cell nucleus model at hand incorporates activity as well as the deformability of the shell and chromatin-lamina linkages from which correlated chromatin motion emerges. The deformability of the lamina shell is also important to be able to work towards understanding nuclear mechanotransduction [76, 77]. For example, there exists an outflux of calcium ions from a cell nucleus above some critical strain deforming the cell nucleus that, in turn, ultimately affect the acto-myosin contractility of the cell [78]. Rigid shell model cell nuclei make it difficult to quantify such effects. However, as we see now such effects may be key to linking changes in chromatin organization to tissue organization.

## IV. A MULTISCALE HYPOTHESIS INSPIRED BY EXPERIMENTS

Even before the onset of neurogenesis, the human forebrain, consisting of precursor cells known as neuroepithelial (NE) cells, is larger than other mammals [79]. It has, therefore, been long hypothesized that that differences in these NE cells may result in expansion of the neocortical primordium [80, 81]. The expansion begins as tangential expansion and then becomes radial as asymmetric NE cell division emerges with one daughter radial glial (RG) cell (and the other daughter a NE cell) [82]. The RG cells do not inherit epithelial features of NE cells and are rather elongated and, presumably, provide patterning for neurons. Since it is difficult to explore this hypothesis in humans and apes, recent experiments study human and ape brain organoids derived from induced pluripotent stem cells (iPSCs) [1]. Intriguingly, human-derived brain organoids exhibit larger surface area than their ape-derived counterparts [1]. In studying the NE-RG cell transition in such brain organoids, an intermediate cell morphology was discovered and named transitioning NE (tNE) cells. In tNE cells, cell shape changes occur prior to the change in cell identity. This intermediate cell morphology is delayed in human brain organoids in comparison with ape brain organoids. Since the delay in tNE formation postpones the transition from tangential-to-radial expansion, this delay, combined with a shorter cell cycle for human progenitor cells, leads to a larger progenitor pool and, thus, typically larger human-derived brain organoids.

To understand the molecular mechanisms behind this delay in tNE formation in human NE cells, time-resolved sequencing analysis helped to identify differential expression in the zinc-finger transcription factor ZEB2 and, so, a potential driver of tNE cell formation [1]. ZEB2 as a driver was then tested in mutant ZEB2+/-brain organoids as well as controlling ZEB2 expression such that the human-derived and ape-derived brain organoids achieve a similar size and morphology with, for example, the addition of doxycycline to induce ZEB2 expression at earlier stages. Additional treatments validated this hypothesis.

Given such findings, we now ask how does ZEB2, and potentially other players, regulate the delay in the NE-RG cell transition? To answer this question requires understanding of what lies in a cell nucleus. Transcription factors are proteins that control the rate of transcription of genetic information. Of course, these proteins themselves need to be made and so we must understand what controls their own expression rates. While there are a number of pathways regulating transcription given that the NE-RG transition is dominated by cells elongating and so changing shape, we are going to pursue a means of regulation that is mechanical in nature—-mechanical in that some initial cell shape change can potentially induce additional cell shape changes with the aid of transcription factors upregulating or downregulating the transcription of particular proteins.

What do we mean by a mechanical means of regulating transcription? Let us consider human iPSCs. Genetic information is stored in the cell nucleus and when combined with histones, forms chromatin. Chromatin is spatially and temporarily organized within the cell nucleus. While a difference in genetic sequence between, say, a chimpanzee and a human, is rather small—-approximately about 1.2% [83]—-perhaps even this rather small difference in genetic sequence translates into potentially larger differences in spatial organization of the genome inside a cell nucleus. Incidentally, Hi-C maps of human versus chimpanzee stems demonstrate differences [84]. Such differences in spatial organization of the genome can potentially translate into differences in gene expression dynamics, such as ZEB2. Moreover, the spatial organization of chromatin can be modified by a change in the shape of a nucleus, which is often due to a change in the shape of the cell with cell nuclear shape often mimicking cell shape [85, 86]. As evidence for this, Golloshi, *et al*., study chromosome organization before and after melanoma cells travel through 12 micron and 5 micron constrictions to find compartment switching between euchromatin and heterochromatin, among other differences, when performing the Hi-C analysis [87].

As the NE cells divide, given the brain organoid is developing in a confined environment, we hypothesize that the additional cells generate compression on, say, a cell of focus. As the compression increases, there is presumably a slight change in cell shape, which may result in a change in nuclear shape, which then may result in a change in the spatial organization of the chromatin. For instance, a slight compression in a particular direction (and hence elongation of the nucleus in the direction perpendicular to the compression), may open up a chromatin region to facilitate/enhance ZEB2 expression. With this enhancement, presumably ZEB2 is able to take on additional functionality.

*We hypothesize that the amount of compression and/or compression rate required to modify the chromatin organization associated with ZEB2 expression varies from human iPSCs to ape iPSCs. More precisely, a higher amount of compression is needed for human iPSCs as compared to gorilla- or chimp-derived IPSCs*.

Once ZEB2 expression increases, there are multiple downstream effects that can impact cell shape. For instance, ZEB2 is a regulator of SMAD signaling that can affect the production of cell-cell junction proteins [88]. Should the upregulation of ZEB2 lead to fewer cell-cell junction proteins at the apical side, then the cells are able to more readily contract at the apical side. Moreover, the actin-binding protein SHROOM3 helps strengthen the stress fibers oriented in such a way to facilitate constriction [89]. Manipulation of SMAD signalling resulted in influencing the onset of tNE morphology, while treatment with LPA countered the apical constriction [1]. In addition to diminishing the strength cell-cell junctions, enhancing apical constriction, microtubule organization may also be affected. It is known that ZEB2 regulates the production of the microtubule-centrosome binding protein Ninein [90], which may help guide the cell fate transition towards a RG cell given that microtubules shape RG morphology [91]. Moreover, Fouani, *et al*. find that ZEB1 switches from being a transcription factor to a microtubule-associated protein during mitosis [92]. And while they do not find the same phenomenon for ZEB2, the multi-functionality for this class of proteins is rather intriguing [93]. In any event, given the downstream changes to cell adhesion and cell cytoskeletal organization to alter the cellular forces at play, the cell shape transition to more elongated cells drives radial-like expansion of the brain organoid.

Our chromatin-reorganization-due-to-cell-compression hypothesis is readily testable using Hi-C at the single iPSC level to determine at what amount of compressive strain does, or does not, alter the chromatin organization pertaining to ZEB2 expression. At the brain organoid level, the mechanical perturbations are self-generated, if you will, by the cells and the influence of the environment in which the brain organoid is embedded. The more cell nuclei become deformed, the more we need their explicit description in cellular-based models. Note that we would like to go beyond the typical biochemical signaling pathways through cell-cell junctions, focal adhesions, or YAP/TAZ by which others have studied nuclear mechanotransduction [94] to explore directly chromatin organization.

Experimental tests of this multi-scale phenomenon will either validate, or not validate, the hypothesis. Here, we ask the question: How can we build minimal, multiscale models to computationally generate such hypotheses prior to performing gene expression experiments such that the modeling informs the experiments as opposed to experiments informing the modeling? We argue that several of the multi-scale pieces are coming into focus to allow us to more readily connect genetic-scale processes to tissue-scale processes, though there is still much to do. The pieces that are coming into focus are cellular-based models for the structure of organoids as well as structural models of deformable cell nuclei containing chromatin.

## V. COMPUTATIONAL TESTING OF THE HYPOTHESIS—VERTEX MODEL WITH EMBEDDED CELL NUCLEI

Now that we have reviewed recent results concerning two of the major players in the process—-cells and cell nuclei, let us now envision how we can embed one into the other. Earlier work has embedded nuclei in single cells in two-dimensions to understand cell motility in confinement [95]. In that work, the cell nuclear cortex is connected to the cell cortex via springs modeling the remainder of the cellular cytoskeleton, including vimentin, beyond the cell cortex. See Figure 2(b). The model is, therefore, more detailed than prior minimal models, while still remaining foundational in that it reveals a new cell polarity mechanism regulated by vimentin. More specifically, the model predicts that (1) cell speed increases with decreasing vimentin, (2) the loss of vimentin increases nuclear deformation and alters nuclear positioning in the cell, and (3) a new polarity mechanism coupling cell directional motion with vimentin via cytoskeletal strength and nuclear positioning thereby providing a mechanism for the abnormally persistent motion of vimentin-null cells, as observed in experiments.

Given that a direct embedding of cell nuclei of every cell using springs, as was done in the two-dimensional confined cell motility study discussed above is computationally costly, we will do something simpler. We will instead make the simplifying assumption that nuclear shape tracks cell shape such that any changes in cell shape result in changes in nuclear shape. For instance, should a cell become elongated, its cell nucleus will also become elongated with the same strain. We will address going beyond this simplifying assumption in the discussion.

To begin to computationally test the notion that chromatin organization can change in response to tissue structure in development, we start with a structure that is reminiscent of a brain organoid structure after several days in development. We will begin with a cortex-lumen structure, which is a band of cells surrounding a fluid-filled core, otherwise known as a lumen. Of course, over the course of several days, the cells in the brain organoid have divided. While incorporating cell division in the three-dimensional vertex model is currently in progress, we mimic an effect of cell division by compressing the organoid in a particular direction. In the confined *in vitro* system, dividing cells become compressed in various directions as they divide. We will do the same, though assuming a particular direction of compression that is applied locally. We do this for organoids with a target shape index of *s*_0_ = 5.6. See Figure 6. We fit each cell to a minimal volume ellipsoid and determine its long axis. In Figures 6(a) and (b), the cell colored red denotes the cell with the largest change in strain along its long axis. In Figures 6(c) and (d), we plot the probability distribution of the aspect ratio of cells before and after the deformation. As a result of the deformation, the probability distribution becomes narrower. Note that the organoid is not confined in the direction orthogonal to the deformation and so not all cells will be more elongated. Since strain is what one can control in the simulations, as opposed to shape, we also plot the cellular strain as a function of simulation time for the cell in red in Figures 6(a) and (b). We will use this information to study how cell nuclear shape changes.

**FIG. 6.**
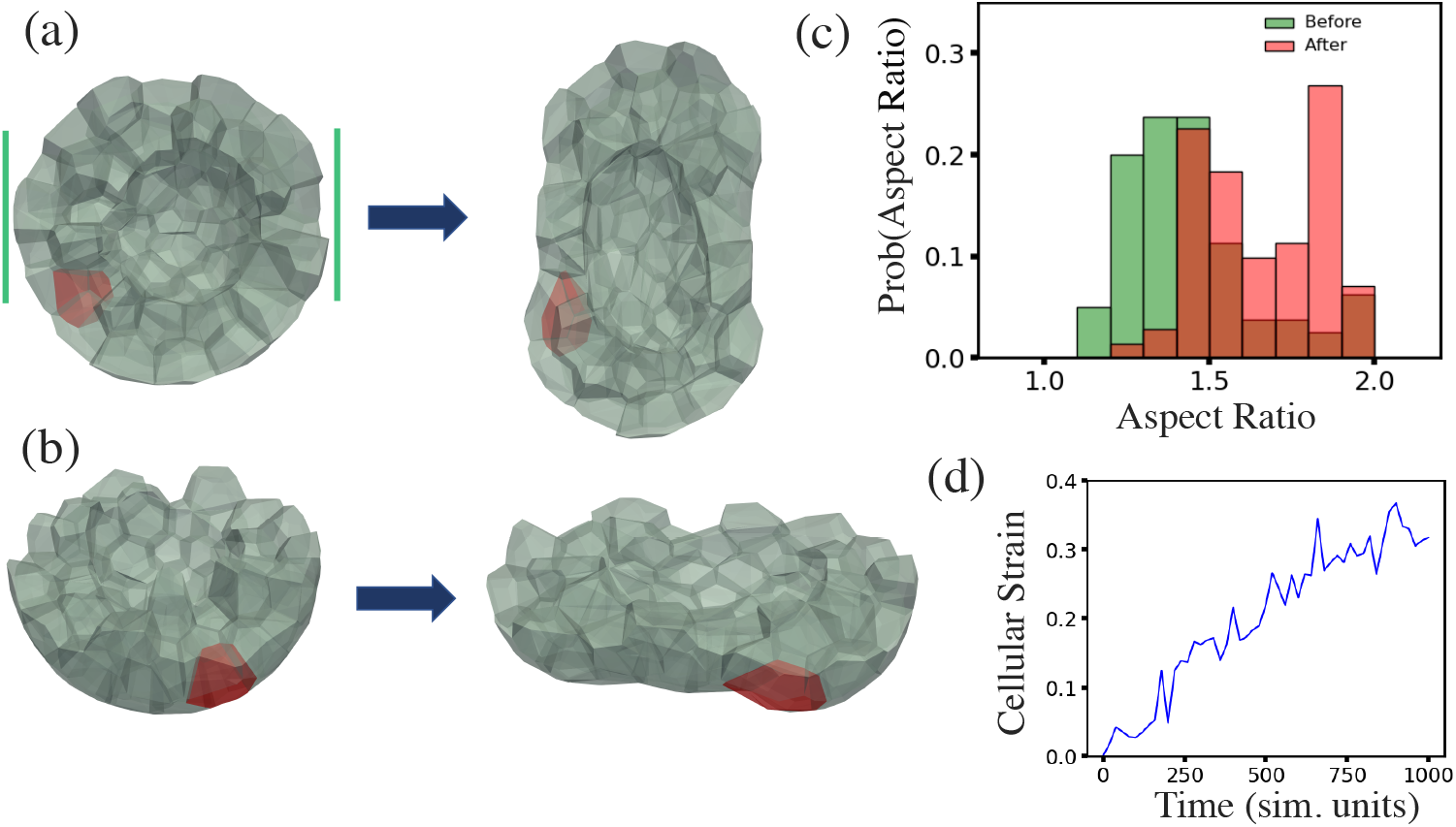
Localized compression a model brain organoid containing a lumen: (a) Top view of before and after compression a fluid-like brain organoid with *s*_0_ = 5.6 and cut in half to better expose the interior. The compression occurs in the region denoted by the vertical green lines. (b) Side view of (a). (c) Probability distribution of the aspect ratio of each cell before and after the local compression. (d) Plot of the cell undergoing the largest change in strain as the organoid undergoes compression. The corresponding cell is labelled in red in (a).

Now that we have measured cell strain changes, which is related to cell shape, in response to tissue-scale compression, we invoke a simplifying assumption to proceed at the next smallest length scale. Our simplifying assumption is that cell nuclear shape tracks cell shape. We can now apply mechanical deformations to our deformable cell nuclei that track cell deformations and ask how does the chromatin organization change in response. Granted, our model for chromatin is indeed a coarse-grained one in that it does not contain a detailed chromatin model with multiple pairs of chromosomes, for example. However, we can ask questions for several example perturbations, such as for a given number of chromatin crosslinks and linkages, by how much does the chromatin chain locally displace in the presence of uniaxial compression? And should the number of crosslinks or linkages be perturbed, by how much does the local displacements change? Do local displacement changes track with the local perturbations? We can indeed still learn things from coarse-grained simulations in terms of generic trends.

Given the amount of deformation in the maximally deformed cell as a function of time, we next apply that same deformation rate to a model cell nucleus. We can then measure the displacement of the chromatin monomers. We can do so for the same initial chromatin configuration, though changing the number of chromatin crosslinks and linkages. For a larger number of chromatin crosslinks, one expects smaller displacements as compared with a smaller number of chromatin crosslinks. However, perhaps even perturbing the number of chromatin crosslinks (for the same initial chain configuration) will lead to pockets of differences in displacements in the chromatin. Such pockets could be candidates for changes in genetic expression, even within this minimal model. For now, we have turned off activity, which could be yet another generator of changes in gene expression for two genetically very similar genomes.

Figure 7 demonstrates our chromatin displacement results. Figure 7(a) shows snapshots from uni-axial compression of the lamina shell by two parallel plates moving at constant speed towards each other and exerting force on the lamina shell, but not on the chromatin. In Figures 7(b) and 7(c) we show chromatin displacement fields for *N*_*C*_ = 2500 and *N*_*L*_ = 400 and *N*_*C*_ = *N*_*L*_ = 0 respectively. We observe smaller displacements for the more crosslinked chromatin. In Figure 7(d), we plot the probability density function of the magnitude of the displacements for the two cases in Figures 7(b) and 7(c). We also include the probability density function for 10 fewer chromatin crosslinks, for the same initial configuration, and find small differences in the distribution from the unperturbed case.

**FIG. 7.**
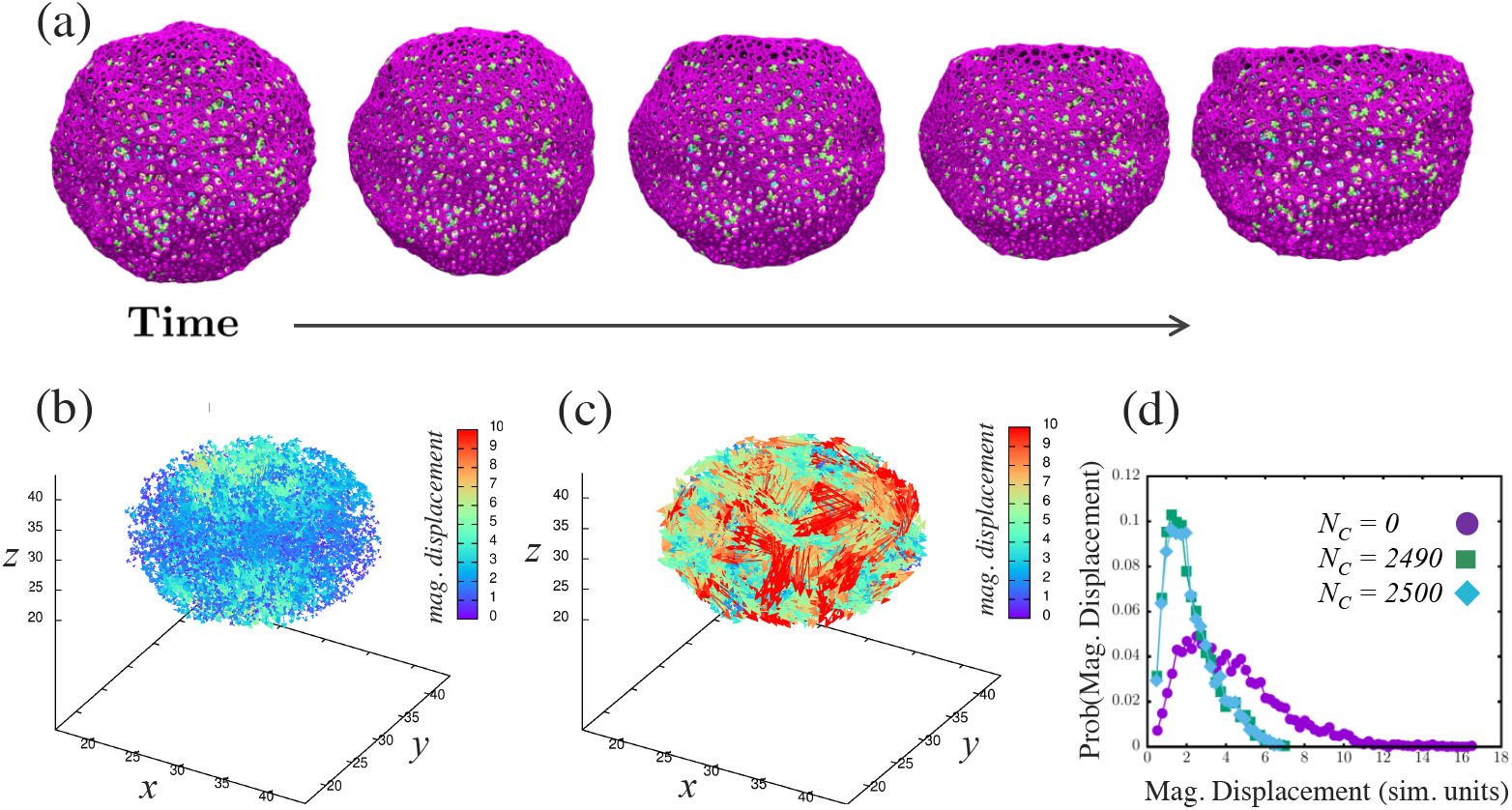
Compressing a model cell nucleus consisting of a lamina shell containing chromatin: (a) Snapshots of the model cell nucleus being compressed for number of chromatin crosslinks *N*_*C*_ = 2500 and number of linkages *N*_*L*_ = 400. The strain rate is the same as in Figure 6(d). (b) Plot of the chromatin displacement field for (a). (c) Plot of the chromatin displacement field for *N*_*C*_ = *N*_*L*_ = 0 for comparison. (d) Probability distribution of the magnitude for the chromatin displacement field comparing (b) with (c) and also for *N*_*C*_ = 2490, i.e., a small change from the configuration in (a).

We can look for spatial correlations in the differences in displacements between the control case, if you will, and the perturbed case with slightly fewer crosslinks. When looking at the spatial map of the differences in displacements (Figure 8), we do find pockets of larger differences in displacements with such regions being candidates for differences in gene expression with the assumption that changes in chromatin configuration can potentially influence genetic regulatory networks at the base pair level. Those pockets become more pronounced the larger the threshold magnitude of displacements. Therefore, it is a worthwhile endeavor to compare differences in displacements from a particular reference state, which could be genetically-close relative, in response to mechanical perturbations. Note that we have only considered one type of perturbation from the reference state here. Other types of perturbations can be explored.

**FIG. 8.**
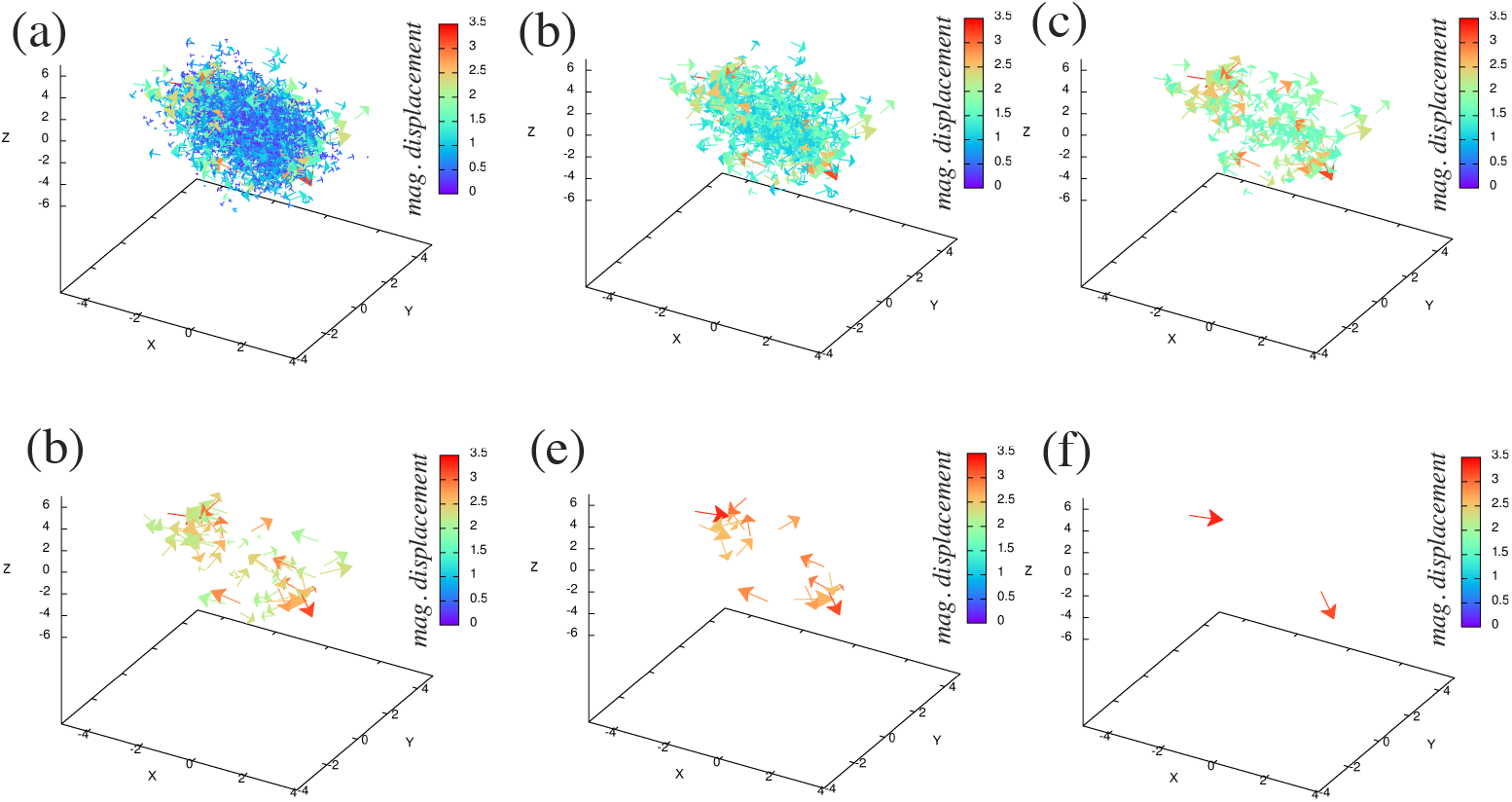
Comparing differences in chromatin displacement fields between the unperturbed and perturbed chromatin configuration with 0.4% change in the number of chromatin crosslinkers: (a)-(f) The threshold magnitude of the difference in chromatin displacements between *N*_*C*_ = 2500 and *N*_*C*_ = 2490 increases from top left to bottom right. By increasing the threshold magnitude, one can more readily identify regions of differences. These regions of differences do not necessarily correspondence to the differences in locations of the chromatin crosslinkers.

If chromatin were a purely a liquid on both short and long spatial scales and time scales, such a comparison between a reference state and a perturbed state would not provide much insight. However, experiments demonstrate that the presence of chromatin crosslinks establish the need for a more intricate rheology of chromatin such that it can act elastically over some time and length scales [96–98]. For instance, mechanical measurements of stretched cell nuclei can be readily explained with chromatin crosslinks [96]. And recent isotropic swelling of cell nuclei reveal very reproducible chromatin configurations before and after the swelling [98]. Other experiments indicate that a Burger’s model fits micropipette aspiration data [99]. Moreover, linkages in terms of LADs between chromatin and the lamina shell also potentially provide some elasticity over shorter time scales [23]. Presumably, linkages dominate closer to the periphery than in the bulk. So while the jury is still out on chromatin rheology, we hope that future chromatin organization studies between perturbed and unperturbed cases, corresponding to human and genetically-close relatives, will become a staple of multiscale structural analysis.

## VI. DISCUSSION

We posit a testable, multi-scale hypothesis for a difference in brain organoid structures derived from human-derived pluripotent stem cells and chimpanzee-derived pluripotent stem cells during the first ten days of development. The hypothesis involves, ultimately, mechanical perturbations of cell nuclei with human-derived pluripotent stem cells demonstrating a different critical strain for particular regions of chromatin organization as compared to chimpanzee-derived pluripotent stem cells relevant to changes in gene expression of the ZEB2 transcription factor that can ultimately impact cell shape by way of decreasing apical cell adhesion and increasing apical cell constriction, as shown previously [1].

While experimental confirmation awaits, we ask what insights can computational modeling provide in terms of building such testable, multi-scale hypotheses. To develop such insights, we argue that using cell-based, computational modeling in terms of, say, a three-dimensional vertex model as presented in Ref. [22] is a reasonable starting point for an organoid, more generally, in the early stages of development. Such models can ultimately provide accurate descriptions of cell shape. We also make the simplifying assumption that nuclear shape tracks cell shape. Now that there is a coarse-grained mechanical model for a cell nucleus that allows for deformability and contains chromatin that recapitulates both the mechanics and correlated chromatin motion [23, 69], we can begin to study, at some level, chromatin reorganization. Many other chromatin-based models of do not allow for nuclear deformability and so are not able to capture such effects [48]. In addition, there is experimental evidence for feedback between cell shape and nuclear shape, with the nucleus releasing calcium should it be compressed above a critical strain [78]. We have not yet explored such feedback here but it certainly unlocks more possibilities in the sense that the cell nucleus shape does not simply track cell shape with perhaps the release of calcium working to preserve the chromatin organization in the presence of particular mechanical perturbations.

While cellular-based models are becoming more predictive in terms of cell shape, what about cell nucleus models? Here, we reviewed a base, mechanical model of cell nucleus, which is a coarse-grained model. Since we are ultimately after a predictive model for changes in chromatin configuration as a function of mechanical and chemical perturbations, the cell nucleus model will require more detail such as heterochromatin versus euchromatin and liquid-liquid phase separation of chromatin crosslinkers [100] and more accurate motor representation to work towards predictive Hi-C in the presence of mechanical perturbations. For instance, condensin II appears to determine genome architecture across species [101]. Efforts are already underway via HiCRes and HiCReg that are rooted in libraries but do not appear to focus on nuclear shape [102]. Moreover, cell division plays an important role and so understanding how chromatin reorganization during cell division is important as well. Finally, while our focus here has been on the downstream effects of ZEB2 one certainly anticipates other proteins to be involved given the collective genomic landscape.

As experimental scientists are able to obtain more detailed information at multiple scales in living systems, it behooves the non-experimental scientists to be able to stitch the scales together not just retroactivel,y but proactively, to be able to better understand the design principles of life. In other words, connecting the dots between genes and tissues theoretically is becoming increasingly within our reach. While here we focused on a multiscale hypothesis for the structure of brain organoids, one can obviously think more broadly to organoids and tissues in general. Of course, certain types of questions do not require as detailed an explicit framework from genes to chromatin to cell nuclei to cells to tissues and so we continue to work on answering them, though not losing sight of the more detailed, multi-scale, computational modeling road that we are just beginning to travel. We hope that many will join us along the way.

J.M.S. would like to acknowledge discussion with Orly Reiner, Amnon Buxboim, Ken Kosik, Ahmad Borzou, Tjitse van der Molen, Alison Patteson, James Li, Alex Joyner, and Ed Banigan. J.M.S. acknowledges financial support from NSF-DMR-CMMT 1832002 and NSFPHY-PoLS 2014192 and from an Isaac Newton Award for Transformative Ideas during the COVID19 Pandemic from the DoD. M.A.L. is supported by the Medical Research Council (MC_UP_1201/9) and the European Research Council (ERC STG 757710).

## Notes

### Competing Interest Statement

The authors have declared no competing interest.

